# Sign of selection on mutation rate modifiers depends on population size

**DOI:** 10.1101/157131

**Authors:** Yevgeniy Raynes, C. Scott Wylie, Paul D. Sniegowski, Daniel M. Weinreich

## Abstract

The influence of population size *(N)* on natural selection acting on alleles that affect fitness has been understood for over half a century^1^. As *N* declines, genetic drift overwhelms selection and alleles with direct fitness effects are rendered neutral. Often, though, alleles experience so called indirect selection, meaning they affect not the fitness of an individual but the fitness distribution of its offspring. Some of the best studied examples of indirect selection include alleles that modify aspects of the genetic system such as recombination^2^ and mutation^3^ rates. Here we use analytics, simulations and experimental populations of *S. cerevisiae* to show that modifiers that increase the genomic mutation rate (mutators) are favored by indirect selection in large populations but become disfavored as *N* declines. This surprising phenomenon of sign inversion in selective effect demonstrates that indirect selection on a mutator exhibits a qualitatively novel dependence on *N*. Sign inversion may help understand the relatively sporadic distribution of mutators in nature despite their frequent emergence in laboratory populations. More generally, sign inversion may be broadly applicable to other instances of indirect selection, suggesting a previously unappreciated but critical role of population size in evolution.

## Main Text

Genomic mutation rates are affected by DNA replication and repair enzymes. Allelic variants of these enzymes that raise the genomic mutation rate are known as mutators. Even when they have no direct effect on fitness, mutators can experience indirect selection via genomic associations with fitness-affecting mutations^3^. Most of these mutations are expected to be deleterious^4^.Nevertheless, theoretical models predict that mutators should be favored in non-recombining populations because they can hitchhike^5^ with occasional, beneficial mutations^6-8^. Consistent with these predictions, mutators have repeatedly spontaneously emerged and achieved fixation in microbial evolution experiments^9-12^.

Notably, most theoretical and empirical studies of mutators have focused on large populations. Building on earlier studies^8,13-16^, we hypothesized that in sufficiently small populations (i.e., not susceptible to interference from competing lineages) mutator alleles can fix by only one of two classically understood processes. Namely, a mutator can fix if genetic drift happens to overcome its increased deleterious mutational load. Alternatively, the mutator may produce a beneficial mutation that escapes drift and sweeps to fixation, taking the mutator with it. Algebraically, we thus approximate the fixation probability of a mutator as

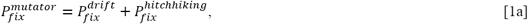

where

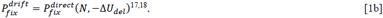

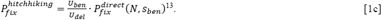

Here, *U_ben_* and *U_del_* are the beneficial and deleterious mutation rates respectively, *ΔU_del_* is the difference in *U_del_* between mutator and non-mutator, *s_ben_* is the selection coefficient of a beneficial mutation and

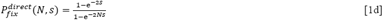

from ^1^ (Methods). As Figure 1 shows, this approximation (now normalized by the fixation probability of a neutral allele, 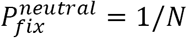 is closely borne out in stochastic Wright-Fisher^19^ simulations of asexual populations.

**Figure 1:**
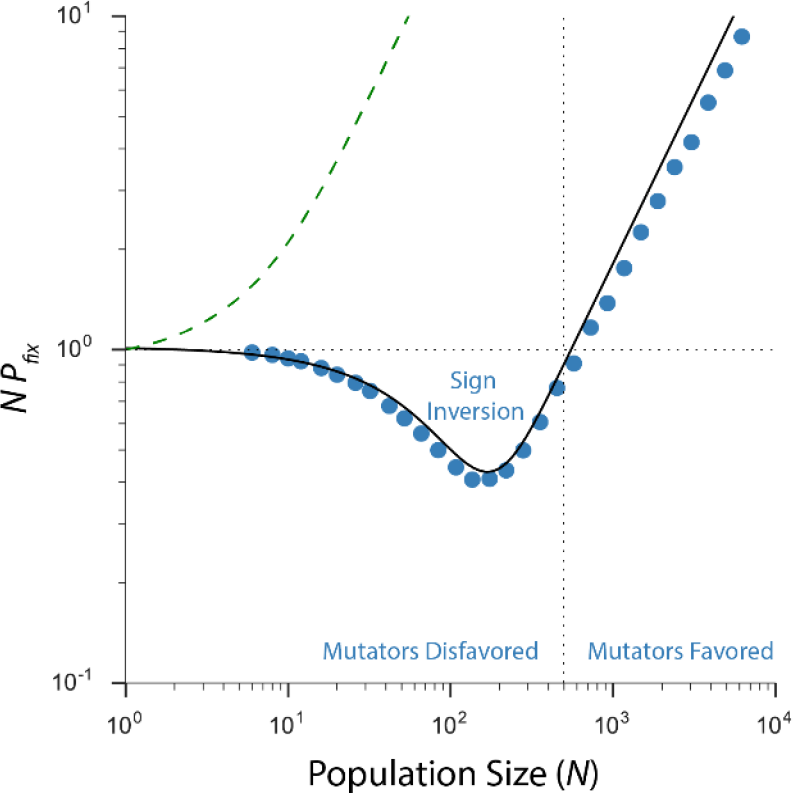
Population size *(N)* determines the sign of indirect selection on mutators. Sign inversion for mutators occurs when their normalized fixation probability (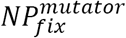) crosses the neutral expectation (horizontal dotted line). Mutators fare better than neutral alleles above 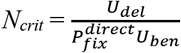 (vertical line), but worse below. Solid black line: heuristic treatment (see text). Blue dots: stochastic simulations, 10^6^ replicates. In contrast, the normalized fixation probability of a directly beneficial mutation (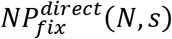, green dashed line) asymptotically approaches but never crosses the neutral threshold. Parameter values: *U_del_*=10^-4^, *U_ben_*=10^-6^, *s_ben_*=0.1, *s_del_*=-0.1. Mutators mutate 100 × faster than non-mutators.

Figure 1 also illustrates the most surprising behavior of 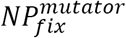:the N-dependent inversion in the sign of selection (“sign inversion”), which occurs when 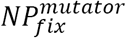 crosses 1, i.e., the neutral expectation: 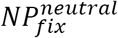. Setting 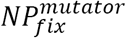 to 1 we find that selection is predicted to favor the mutator above 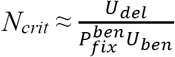 and disfavor it below. This transition occurs because selection against the deleterious mutational load makes mutator fixation by drift (Eq. 1b) very unlikely (generally 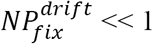) (Extended Data Figure 1). Meanwhile mutator hitchhiking probability (Eq. 1c) is everywhere depressed compared to a directly favored mutation (Eq. 1d) by the requirement to first generate such a mutation. Consequently, unlike 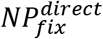, which asymptotically approaches the neutral expectation as *N* declines (Figure 1), 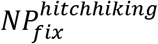 drops below it at *N_crit_* (Extended Data Figure 1). Once this occurs, selection against the deleterious load keeps 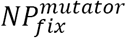 below the neutral expectation until the cost of the load is overpowered by drift (i.e. 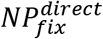 approaches 1).

**Extended Data Fig. 1.**
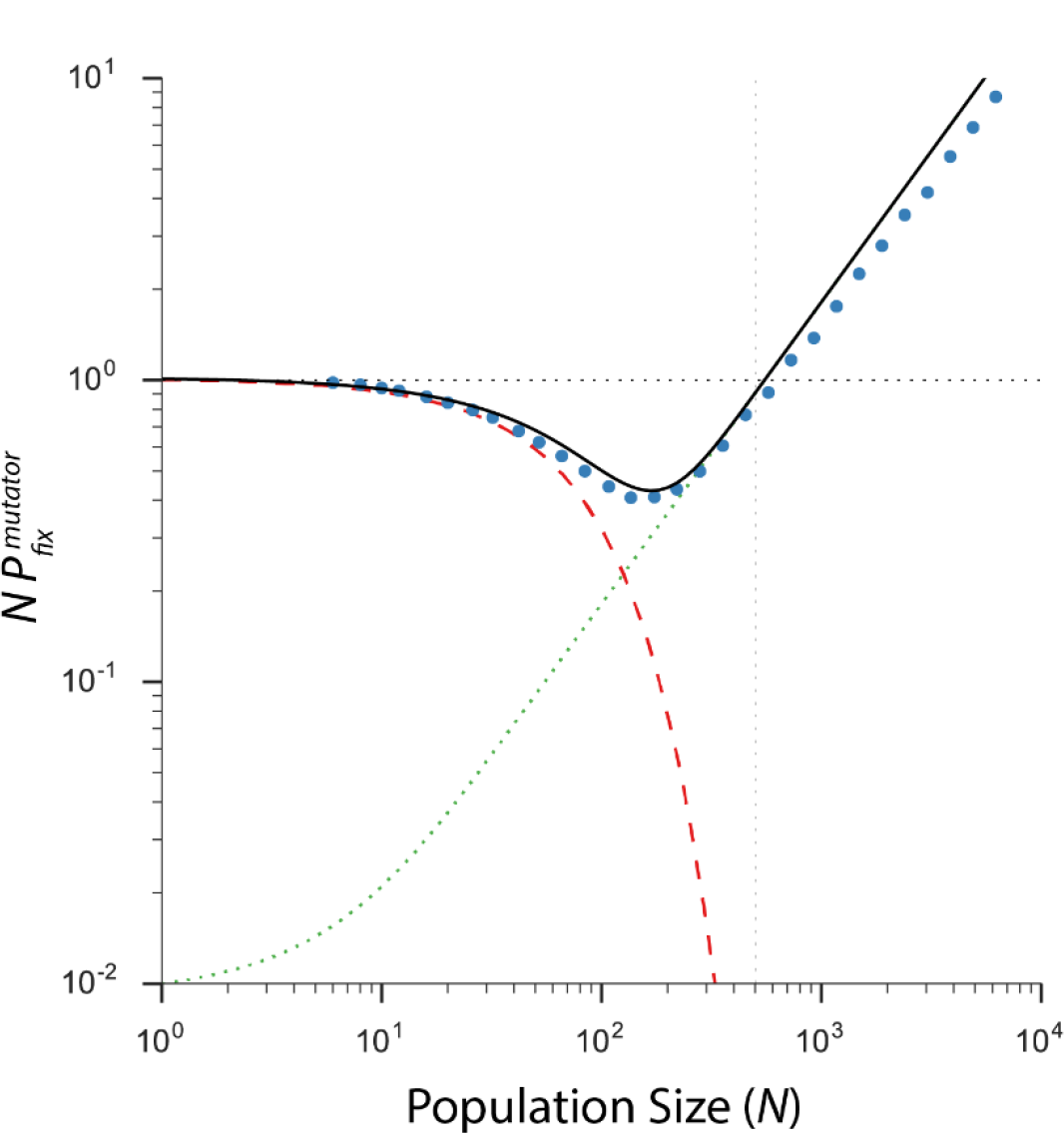
Sign inversion occurs when the scaled fixation probability of a mutator (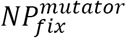) crosses the neutral threshold (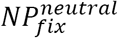 = 1, horizontal dotted line). Analytic expectation (black solid line) is the sum of 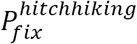 (green dotted line) and 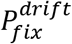 (red dashed line). 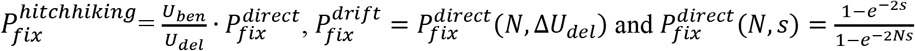 (See Main text and Methods for derivations.) Blue dots: simulation results, 10^6^ replicates. Parameter values: 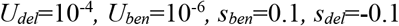. Mutators mutate 100× faster than non-mutators, giving 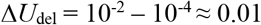 Vertical line: 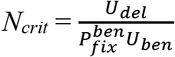

To empirically test the prediction of sign inversion we conducted competitions between mutator and non-mutator strains of *S. cerevisiae*. We initiated competitions at approximately equal frequencies (thereby changing the neutral fixation probability from 1/*N* to ~0.5) and propagated them at three different population sizes by regular dilutions into fresh medium. Following other experimental studies and population genetics theory ^14,20,21^ we manipulated population size by varying the size of the dilution bottleneck. Simulations confirmed that small bottlenecks can also cause sign inversion (Extended Data Figure 3). Based on simulations, we predicted that sign inversion would occur at a bottleneck somewhere between ~50 and ~1000 cells (Extended Data Figure 3).

**Extended Data Fig. 3.**
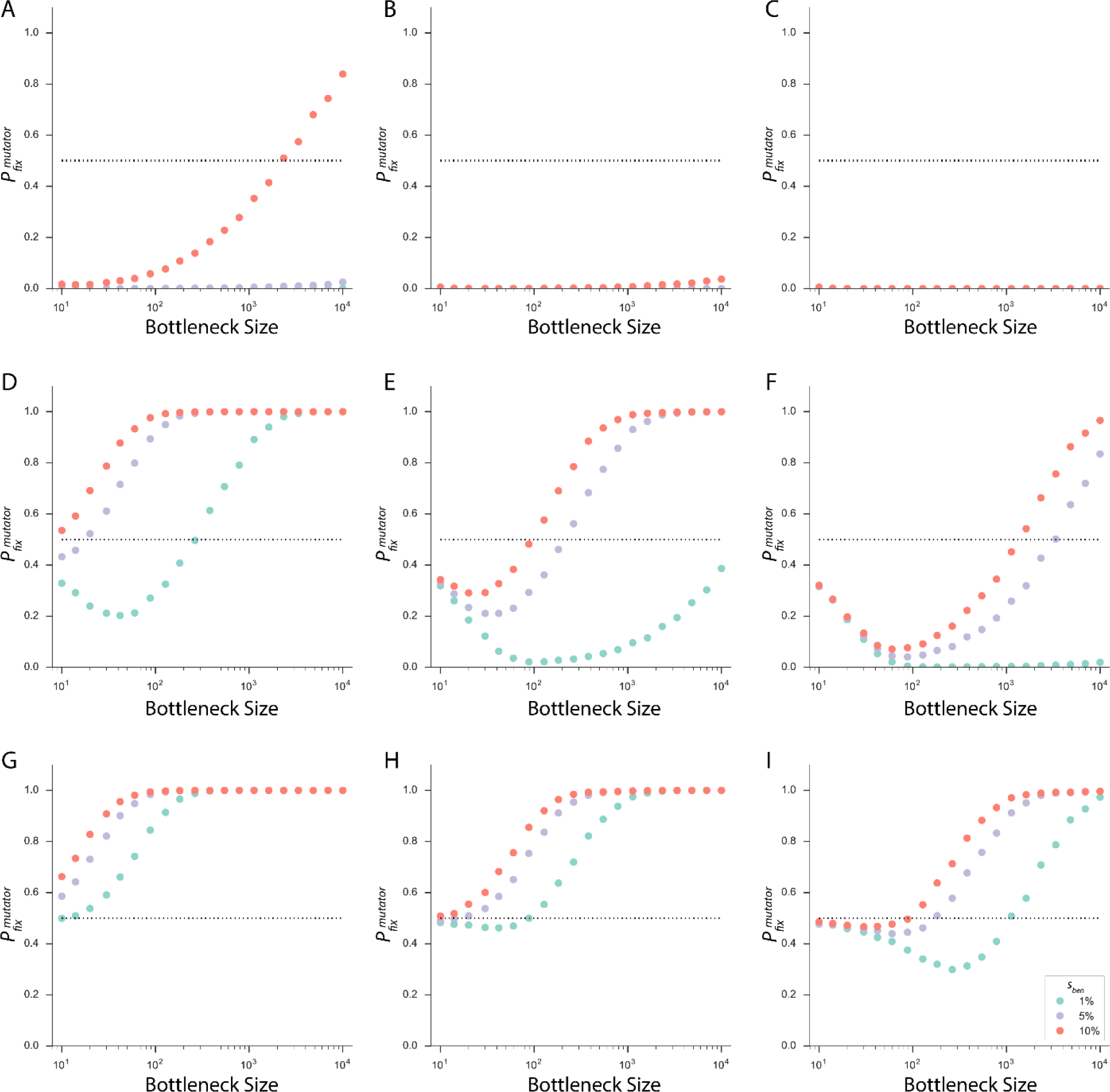
Sign inversion in simulated bottleneck populations. To inform experimental design, we conducted simulations of bottlenecked populations started with the mutators at the frequency of 50%, raising 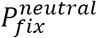 to 50% (note that unlike everywhere else, here 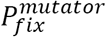 is not normalized by 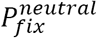). Similarly to simulations in constant-size populations, 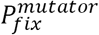 declines below the neutral expectation with decreasing bottleneck size. Simulations were conducted with *N_max_* = 8×10^5^ (Methods) to correspond to the experimental medium. Because the true values of *U_del,_ U_ben,_* and *s_ben_* in our experimental medium are unknown, in simulation we explored values from other experiments (Methods): we varied *U_del_* between 10^-3^ (row 1), 10^-4^ (row 2), and 10^-5^ (row3), and we varied *U_ben_* between 10^-5^ (column 1), 10^-6^ (column 2), and 10^-7^ (column 3). Based on these simulations we estimate that the critical bottleneck size in experimental populations of yeast may be approximately between 50 and 1000 cells. 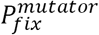 calculated over 125,000 runs.

Figure 2 presents the results of competitions propagated through large bottlenecks of ~8000 cells (2A), intermediate bottlenecks of ~80 cells (2B), and small bottlenecks of ~20 cells (2C). As predicted, mutators were strongly favored in large populations but strongly disfavored in small populations. In large populations mutators won in 162 of the 276 competitions (see Methods) in the first ~250 generations (~58.7%, significantly above the neutral expectation; two-sided, binomial test *p* < 0.01) and lost in only 25 (Figure 2A). In addition, the mean mutator frequency in the 89 unresolved competitions (~0.63±0.03 S.E.M.) was also significantly higher than the starting frequency (two-sided, paired t-test t_df = 88_ = 4.4870, *p* << 0.01), consistent with selection favoring mutators. In intermediate populations, mutators won in 112 of 275 competitions (~40.7%) and lost in 124 (~45.1%) in the first ~225 generations (Figure 2B). The mean mutator frequency in the remaining 39 competitions (~0.49±0.05 S.E.M.) was not significantly different from the mean starting frequency in these populations (two-sided, paired t-test t_df=38_ = 0.1747, p = 0.8622). Selection for mutators was thus clearly weakened by intermediate bottlenecks and it appears that these populations are close to *Ncrit*. Finally, in small populations (Figure 2C), mutators won in 53 of 273 competitions (~19.4%; significantly below the neutral expectation; two-sided, binomial test *p* << 0.01) and lost in 208 (~76.2%) in ~250 generations. The mean mutator frequency in the remaining 12 competitions (~0.43±0.07 S.E.M.) was not significantly different from their starting frequency (two-sided, paired t-test t_df=11_ = 0.8173, p = 0.4311)

**Figure 2:**
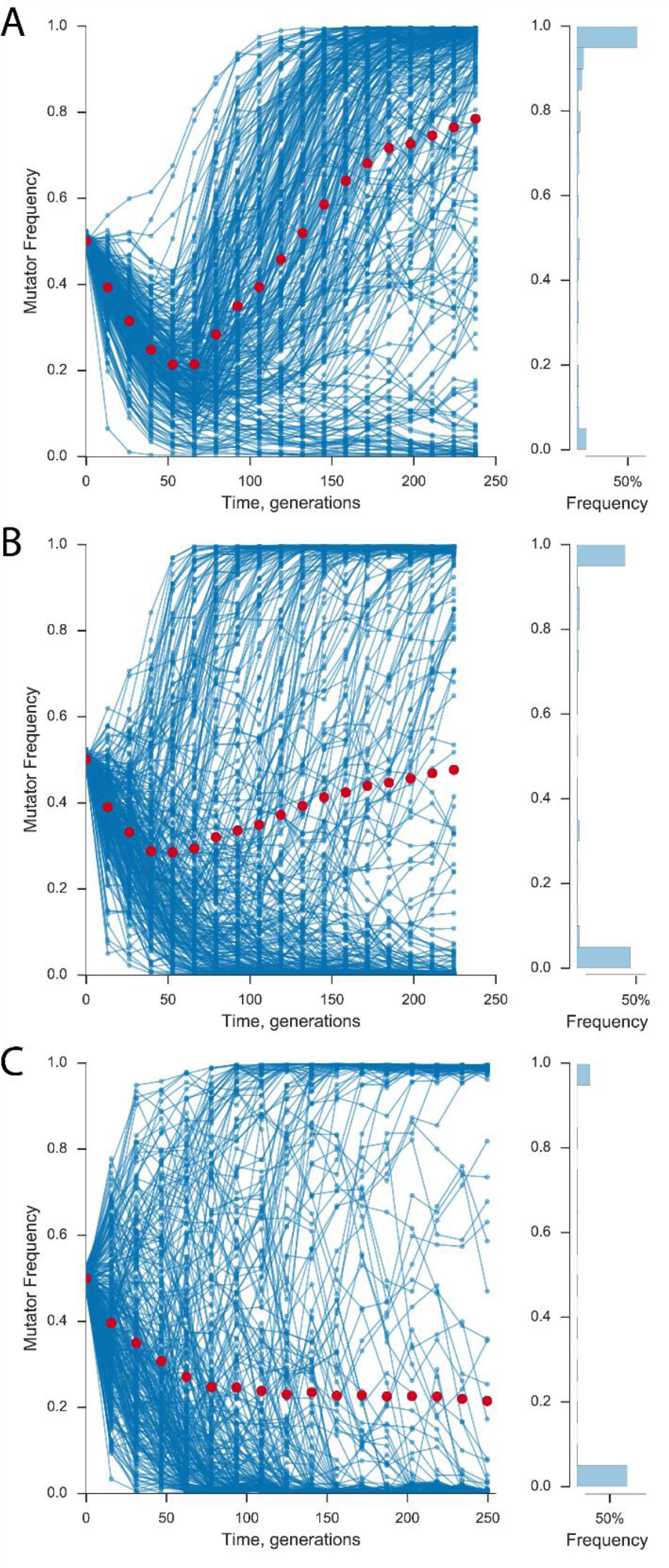
Sign inversion in experimental yeast populations: Mutators are favored in large populations but disfavored in small populations. Left: Mutator dynamics in populations propagated through bottlenecks A) Large (~8000 cells, B) Medium (~80 cells), and C) Small (~20 cells). Blue curves: individual population frequencies; red dots: all population averages. Right: frequency histograms on the last day of propagation.

To confirm the mechanism driving sign inversion in our competitions, we measured mean fitness values of winning mutator and non-mutator populations. Consistent with our expectation that above *N_crit_* mutator fixation is primarily driven by hitchhiking with beneficial mutations, all mutator winners in simulated (Figure 3A) and experimental (Figure 3B) large populations were considerably and significantly fitter than their ancestors (in experiments: two-sided t-tests, all *p* < 0.01, Supplementary Data 2).

**Figure 3:**
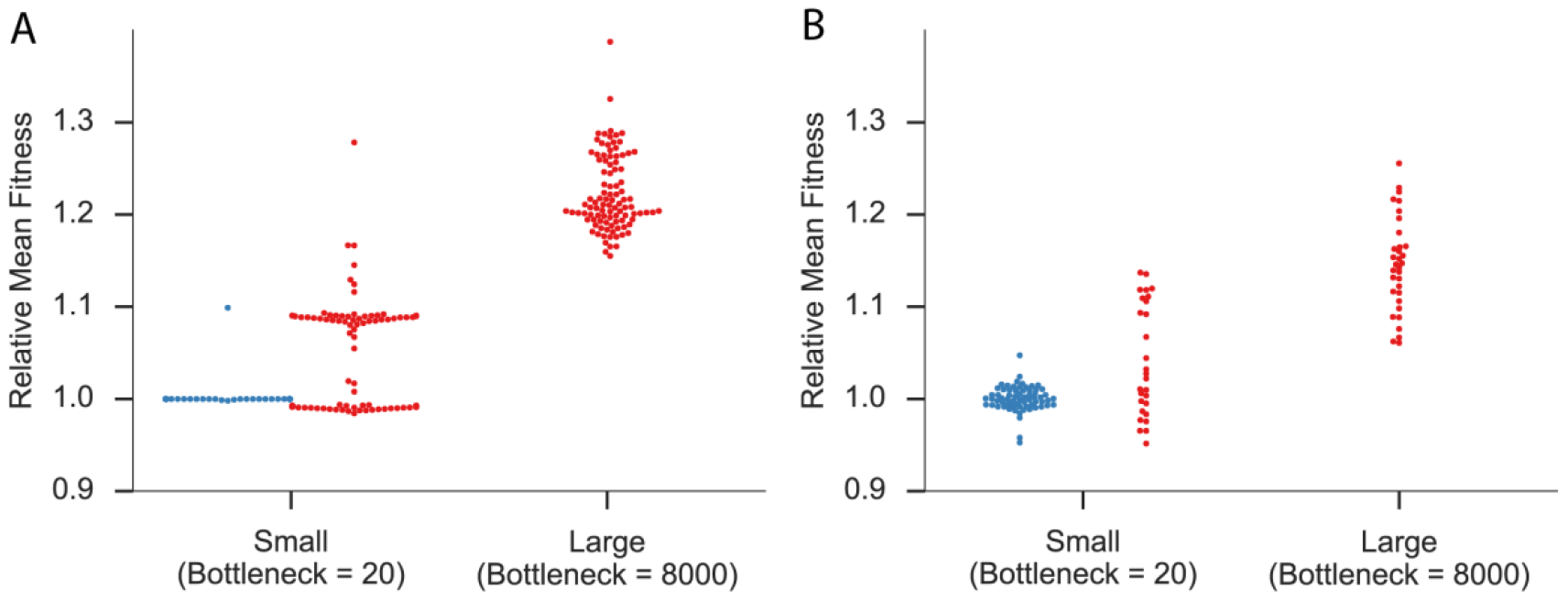
Mutators win large competitions by hitchhiking, non-mutators win small competitions by outlasting the mutators. **A)**Mean fitness of simulated competition winners. We simulated 50/50 competitions propagated through small bottlenecks of 20 cells and large bottlenecks of 8,000 cells. As expected mutators are favored at large 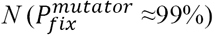 and disfavored at small 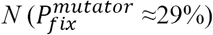. Furthermore, large *N* mutator winners are always more fit than their ancestors. In contrast, only some small *N* mutator winners are more fit than their ancestors. Meanwhile, non-mutator winners of small *N* simulations are almost never more fit than their ancestors. 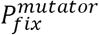averaged over 10^6^ runs. Each point represents population fitness at the end of simulation (a random 100 replicates shown for clarity; note that 99/100 non-mutator winners have a relative fitness of one). Parameter values are the same as in Figure 1. **B)** Mean fitness of experimental competition winners. Mean fitness of large populations in which the mutator had won is always considerably higher than their ancestors, consistent with hitchhiking. In contrast, only some small populations in which mutators had won show evidence of hitchhiking. Mean fitness of small populations in which non-mutators had won is very close to their ancestors, ruling out hitchhiking. Each point represents an average of 5 replicate fitness assays. See Extended Data Figure 4 for 95% CI.

In contrast, below *N_crit_* mutators may fix by either hitchhiking or genetic drift. Correspondingly, in both simulated (Figure 3A) and experimental (Figure 3B) small populations some of the mutator winners were considerably fitter than their ancestors (i.e., fixed by hitchhiking), while some were about as fit or even less fit than their ancestors (i.e., fixed by drift). However, non-mutators are expected to fix not by hitchhiking but by outlasting the mutators lost to selection against the deleterious load. Indeed, non-mutator winners of simulated small populations (Figure 3A) were almost never fitter than their ancestors. Likewise, most non-mutator winners of experimental small populations (Figure 3B) were not significantly fitter than the ancestor while the rest were not fit enough (~1% more fit on average) to have fixed by hitchhiking so quickly (Supplementary Data 2). Thus we reason that their fixation was most likely still driven by outlasting the mutators lost to the deleterious load.

Taken together our results demonstrate that indirect selection favors mutators in large populations but suppresses them in small populations. Understanding the role of *N* in mutation rate evolution helps elucidate where and why mutators are likely to prevail. In particular, it may help explain why mutators often emerge in experimental microbial populations but have been rarely found in nature, except sporadically in cancer cells^22,23^ and pathogenic infections^24,25^. Importantly, whereas natural populations are likely to be large, they are also frequently structured in both time and space (e.g., transmission bottlenecks^26^, biofilms^27^). Our results suggest that population structure may help inhibit mutators in nature by dividing large populations into smaller demes, in which mutators may be more vulnerable to selection against the deleterious load.

More generally, our results dramatically extend our understanding of the influence of *N* on natural selection. It has been long known that small *N* can weaken selection on alleles that directly affect fitness, rendering them neutral. We have shown that *N* can also change the sign of indirect selection on mutators. Intriguingly, the mechanistic basis of sign inversion may not be unique to mutators. Any mechanism that introduces random genetic or phenotypic variation is more likely to negatively affect fitness than to improve it. Thus, other indirectly selected modifiers of variation, such as modifiers of recombination^28^, may also experience sign inversion. As in the case of mutators, these alleles may be unlikely to achieve fixation by drift due to selection against the predominantly deleterious variants they produce. Meanwhile, their probability of fixation via hitchhiking with beneficial variants may be tempered by the low probability of producing such variants. Sign inversion may thus apply broadly to indirect selection, revealing a profound and previously unappreciated^29,30^ role of population size in evolution.

## Acknowledgements

We thank Christopher Graves, Yinghong Lan, Chintan Modi and other members of the Weinreich laboratory for discussion. We thank Meredith Crane and Amanda Jamieson for help with flow cytometry, John Koschwanez and Andrew Murray for sharing yeast strains and Clifford Zeyl for sharing the MSH2 knockout allele. We thank Casey Dunn, James Kellner, and Sohini Ramachandran for comments on the manuscript. The work was supported by National Science Foundation Grant DEB-1556300.

## Author contributions

Designed project: Y.R, C.S.W, P.D.S, and D.M.W. Developed theory: C.S.W, Y.R, P.D.S, and D.M.W. Conducted simulations: Y.R. and C.S.W. Conducted experiments and analyzed data: Y.R. Wrote the paper: Y.R, C.S.W, P.D.S, and D.M.W

## Author information

The authors declare no competing financial interests. Correspondence and requests for materials should be addressed to Y.R.

## Methods

### Analytic Approximation for 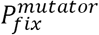

To derive the approximation 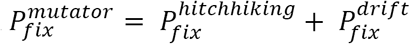, we consider a simple population-genetic model in which mutators can only be enriched by association with beneficial mutations, impeded by association with deleterious mutations, or fix by neutral genetic drift. As in ^13^, we handle new mutations by immediately classifying their long-term fate. We assume that deleterious mutations are destined for extinction, i.e., have a zero fixation probability. Thus a mutator allele can never fix in linkage with a deleterious mutation (Extended Data Figure 2A). In contrast, beneficial mutations fix with probability 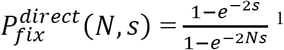. This approximation assumes no clonal interference between contemporaneous beneficial mutations^16,31^ (although this assumption is relaxed in our simulations).

**Extended Data Fig. 2.**
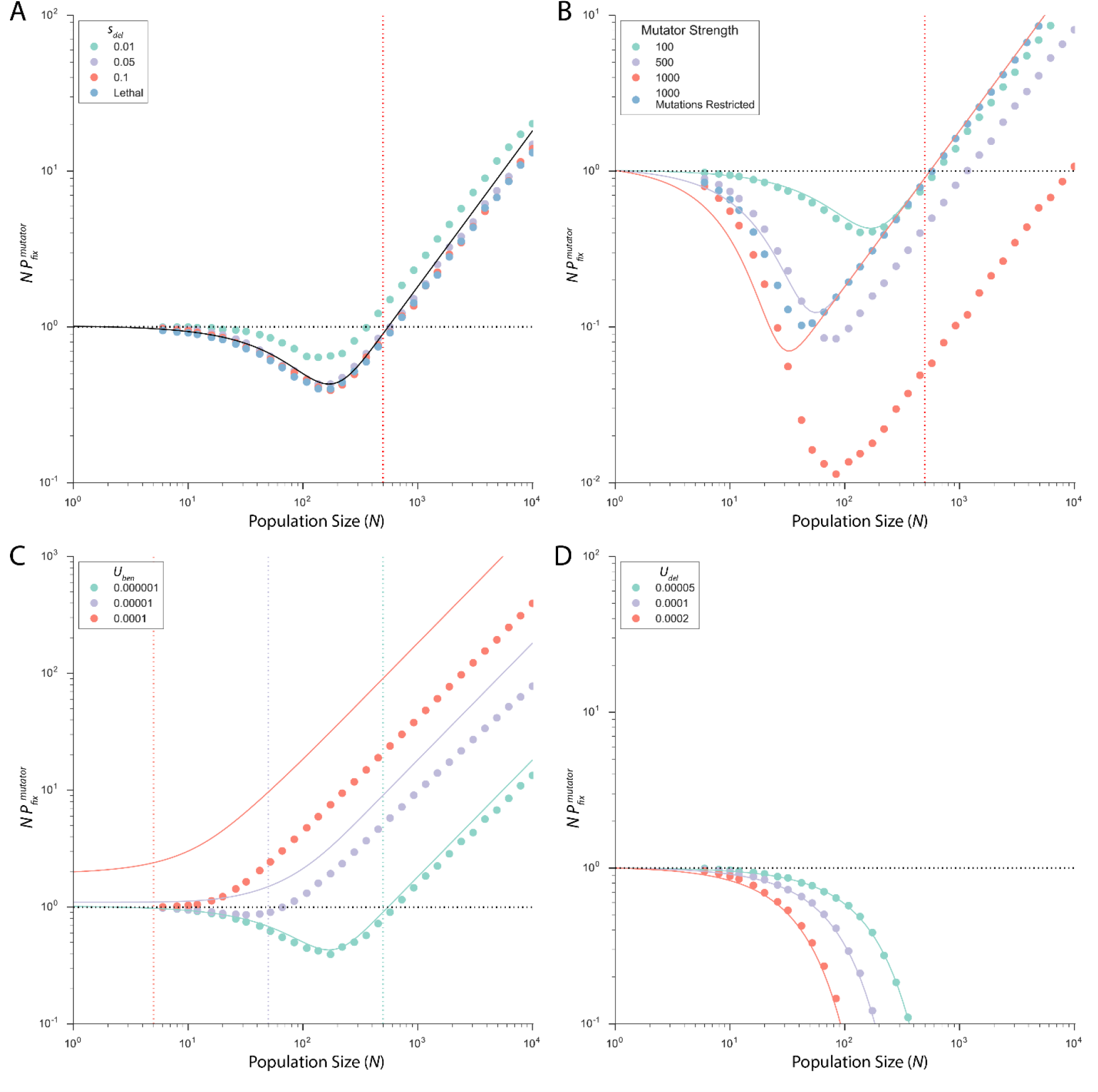
Simplifying assumptions made in the derivation of 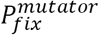. In each panel, vertical dashed lines represent *N_crit,_* horizontal dashed line represents the neutral expectation for 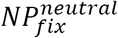, namely 1. Parameter values: *U_del_*=10^-4^, *U_ben_*=10^-6^, *s_ben_*=0.1, *s_del_*=0.1 mutator strength =100-fold, except as noted. a) Model assumes all deleterious mutations are effectively lethal. Simulations show that sign inversion occurs even for deleterious mutations of mild effect and the realized *N_crit_* is almost unaffected. As expected, selection against the deleterious load of mild mutations is weaker than *ΔU_del_* (because populations do not instantaneously achieve mutation-selection balance unless deleterious mutations are lethal). Nevertheless, 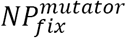in simulation under sufficiently strong deleterious mutations are practically indistinguishable from those under lethal ones. 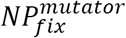 calculated over 10^6^ simulation runs.
b) Model assumes mutator fixation is determined by the first successful beneficial mutation, and not by subsequent beneficial or deleterious mutations. In particular, hitchhiking cannot be spoiled by subsequent deleterious mutations. Simulations confirm that the analytic approximation for 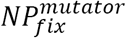 overestimates the realized 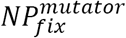 of very strong mutators. These fare worse than predicted because subsequent deleterious mutations spoil some beneficial mutations. This can be seen when new mutations are restricted to only non-mutated backgrounds. Now 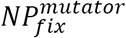 of even very strong mutators is well approximated analytically. 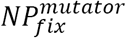 calculated over 5 × 10^6^ runs.
c) Model assumes that deleterious mutations greatly outnumber beneficial ones *(U_del_>>U_ben_)*. Simulations confirm that the analytic approximation for 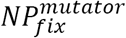 (solid lines) fails to predict realized 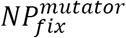 in simulation when *U_ben_*approaches *U_del_*. Although note that our expression for *N_crit_* does not break down in the same way. 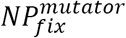 calculated over 10^6^ runs.
d) Model assumes that 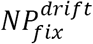 depends on the strength of selection against the deleterious load *~ΔU_del_*. Simulations in the absence of beneficial mutations *(U_ben_* = 0) confirm that 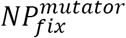 is predicted by 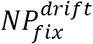(solid lines). 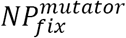 calculated based on 10^6^ runs of the simulation.

We assume that a mutator can fix either by hitchhiking with a successful beneficial mutation or by random genetic drift. The probability of producing a beneficial mutation is the ratio of the beneficial to total mutation rate 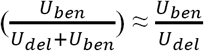, as typically *U_del_>>U_ben_* (Extended Data Figure 2C). 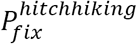 is therefore approximately equal to 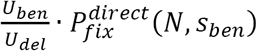. This approximation assumes that sweeping beneficial mutations are unaffected by further mutations in the same background (Extended Data Figure 2B).

In addition to hitchhiking with a beneficial mutation, a mutator allele can fix by random genetic drift, if drift can overpower indirect selection against the excess deleterious load. In the absence of beneficial mutations the disadvantage of a mutator due to the deleterious load is predicted to be *ΔU_del_*^18^ in populations at mutation-selection equilibrium^17^. Assuming, as above, that all deleterious mutations are lethal (i.e. population at the mutation-selection equilibrium), we find 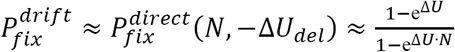 (Extended Data Figure 1). Correspondingly, in the absence of beneficial mutations, 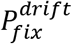 alone predicts 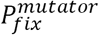 in simulation (Extended Data Figure 2D).

(For a more detailed analytic treatment of 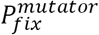 see ^13^.)

### Stochastic simulations

We consider an asexual population evolving in discrete, non-overlapping generations.

The population is composed of genetic lineages, i.e. all individuals that have the same genotype. A genotype is modeled as a vector of 100 loci, including 99 fitness-affecting loci and one mutation rate modifier. The mutation rate locus acts solely on the mutation rate (i.e., it has no intrinsic effect on fitness), and cannot be mutated during simulation. The fitness loci can generate both deleterious and beneficial mutations. We assume fixed fitness effects for all mutations and that they are additive in nature. Fitness is thus calculated as the sum of fitness contributions of all mutated fitness-affecting loci: given x deleterious mutations and *y* beneficial mutations (with selection coefficients of *s_del_* and *s_ben_* respectively) fitness is *w* = 1 - *xs_del_ + ys_ben_*.

Simulations start with *N* individuals divided into two genetic lineages – mutators and non-mutators, all free of fitness-affecting mutations and only differing in their mutation rate. Simulations end when mutators either reach fixation (frequency of 100%) or go extinct (frequency of 0%). Fixation probability is then calculated as the fraction of replicate simulation runs in which mutators fixed. Every generation, the population reproduces according to the Wright-Fisher^19^ model, in which the representation of each genetic lineage in the next generation is drawn from a multinomial distribution with expectation determined by its frequency multiplied by its relative fitness in the previous generation. Unless otherwise stated all simulations were conducted with populations of constant size (N). In simulations of bottlenecked populations, populations double every generation until they reach the maximum population size *N_max_* (=8×10^5^ cells in all simulations; this is the approximate carrying capacity in our experiments). Populations are then reduced to the bottleneck size by sampling every lineage from the population with probability proportional to its frequency.

Upon reproduction, each lineage acquires a Poisson distributed number of mutations, *X* with mean determined by the size of the lineage multiplied by the total per-individual mutation rate *(U_del_* + *U_ben_)*. The counts of beneficial and deleterious mutations are then drawn from a Binomial distribution with *n* = *X* and 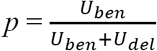. Loci to be mutated are randomly chosen from the non-mutated ones. Simulations were written in Julia 0.5 and are archived as described below. Simulations were conducted with empirically supported parameter values from experimental yeast populations^32-35^: *U_ben_* ~ 10^-6^ - 10^-5^, *S_ben_* ~ 0.02 - 0.1, and *U_del_* ~ 10^-3^ - 10^-5^.

### Strains, media and propagation conditions

Isogenic strains yJHK111and yJHK112 labeled with ymCitrine and ymCherry respectively were generously provided by the laboratory of Andrew Murray and have been previously described ^36^. To engineer mutator strains, we replaced the coding sequence of *MSH2*, a component of the yeast mismatch repair pathway, in both yJHK111 and yJHK112 with a kanamycin resistance knockout cassette as previously described^37,38^. The cassette provides resistance to the antibiotic geneticin (G418) and has been previously shown to have a minimal effect on fitness ^39^ Successful replacement of *MSH2* was confirmed with PCR. Fluctuation tests conducted as previously described^37^ indicated an approximately100-fold increase in the mutator mutation rate toward nystatin (at 4 μM) resistance. Low-glucose minimal medium (6.7g YNB+Nitrogen, 0.2g Glucose per 1L) was used in all experiments to reduce the carrying capacity and lower population size. Medium was supplemented with ampicillin (100μg/mL) and tetracycline (20μg/mL) to prevent bacterial contamination. Populations were propagated in 200μL of medium in wells of standard round-bottom 96-well plates. Plates were sealed in plastic Ziploc bags to prevent evaporation and incubated at 30C with shaking at 1250rpm in a microplate shaker (Multi-Microplate Genie, Scientific Industries, Inc). All populations were periodically frozen in 15% glycerol at -80C.

### Experimental propagation

Preliminary fitness tests as described below indicated a slight but significant advantage of ymCherry over ymCitrine in both the non-mutator background (ymCherry relative fitness = 1.006, t_df = 4_ = 2.9714, *p* = 0.04) and the mutator background (ymCherry relative fitness = 1.013, t_df=4_ = 6.4172, *p* = 0.003). To control for this fitness difference, we employed a blocking design in which half of the competitions were conducted with mutators labeled with ymCherry and half with mutators labeled with ymCitrine. To start, ymCherry (mutator and non-mutator) and ymCitrine (mutator and non-mutator) labeled strains were streaked onto YPD agar plates from frozen stocks. After two days of growth, 288 individual colonies of each of these four genotypes were picked into 200μL of medium and allowed to reach saturation. All populations were then diluted 10,000-fold and allowed to regrow for two more days to physiologically acclimate to the new conditions before competitions started. Once saturated, appropriate mutator and non-mutator cultures were combined at equal volumes into 288 independently founded competitions between ymCherry labeled mutators and ymCitrine labeled non-mutators and 288 independently founded competitions between ymCitrine labeled mutators and ymCherry labeled non-mutators. Mutator frequencies in all populations were then estimated by analyzing about 20,000 cells on the Attune NxT Flow Cytometer (Invitrogen). The 138 populations of each labeling scheme (276 total) closest to 50% mutator frequency were chosen for the experiment. Populations bottlenecked at 8000 and 80 cells were initiated from the same set of 276 populations. Populations bottlenecked at 20 cells were initiated from a new set of 276 populations started in the same way as described above. In the populations used to start the two sets of larger populations, ymCherry labeled mutators were initially at an average frequency of 49.7% ± S.D. 1.17% and ymCitrine labeled mutators were at an average frequency of 50.8% ± S.D. 1.12%. In the populations used to start the 20 cell competitions, ymCherry labeled mutators were initially at an average frequency of 49.9% ± S.D. 1.92% and ymCitrine labeled mutators were at an average frequency of 49.9% ± S.D. 1.87%. Experiments were performed in 96-well plates with four wells per plate filled only with medium to control for cross-contamination.

Large bottleneck populations were propagated by 1:100 daily dilutions into fresh medium, resulting in a bottleneck of about 8000 cells and about ln_2_(100) = 6.6 generations between transfers. Populations bottlenecked at 80 cells were transferred every two days through 1:10,000 dilutions into fresh medium, resulting in about ln_2_(10,000) = 13.2 generations between transfers. Populations bottlenecked at 20 cells were transferred every 2.5 days through 1:40,000 dilutions into fresh medium, resulting in ln_2_(40,000) = 15.3 generations between transfers. One of the populations bottlenecked at 80 cells and three of the populations bottlenecked at 20 cells were lost to accidental extinction during dilution and excluded from the data set. Mutator frequencies were assessed by periodically analyzing about 20,000 cells from each population on the Attune NxT Flow Cytometer.

Competitions were propagated until the sign of indirect selection became clear as assessed by comparing the average frequency and the realized probability of mutator fixation to our neutral expectation (the starting frequency of mutators in our competitions). Note that because the timing of mutator fixation and mutator loss are unknown we did not directly compare realized fixation and loss probabilities after stopping the experiment.

In simulations of bottlenecked populations (Extended Data Figure 3) we observed that the probability of fixation (reaching 100% frequency) for a mutator was almost indistinguishable from the probability of reaching 95% (over 10^6^ replicates): mutators failed to fix after reaching 95% in ~0.1% of simulated small populations and considerably fewer of intermediate (~0.001%) and large (none) populations. Likewise, the probability of extinction in simulations was practically the same as the probability of dropping below 5%: mutators went on to fix after having dropped to below 5% in ~0.08% of small populations, ~0.2% of intermediate populations, and ~0.001% of large populations. In our analysis we thus considered a mutator to have won when it reached a frequency of at least 95% and lost when its frequency dropped to at most 5%.

### Fitness assays

Relative fitness was assayed in short-term competition experiments. Competitors were first inoculated into 200μL cultures in our low-glucose medium and grown to saturation overnight. They were then diluted 1000-fold into the same medium, and allowed to regrow for another day to acclimate to the growth environment. After 24 hours (~10 generations) competitors were combined 50:50 in wells of a 96-well plate and propagated through two 1:100 bottlenecks (two days, 13 generations). The frequency of competitors before and after 13 generations of growth was determined by flow cytometry using the Attune NxT Flow Cytometer. Relative fitnesses were calculated from the change in competitors’ frequencies over the 13 generations of competition using standard population genetics^40^. Each competition was replicated five times.

Winning mutator and non-mutator populations were isolated by sampling experimental (whole) populations in which they had reached a frequency > 99%. Frozen stocks were sampled from the first available time point immediately after winners had reached >99% to minimize the effect of further adaptation after fixation. Winning non-mutator lineages from populations bottlenecked at 20 cells were isolated from populations frozen after about 60 generations (4 transfers) or 120 generations (8 transfers) of propagation. Since mutators were able to fix in only a single population after 60 generations (Figure 2), all mutator winners from our smallest populations were isolated after 120 generations of propagation. Mutator winners of our largest populations were isolated from populations frozen after 192 generations of propagation.

To control for the disadvantage of the higher deleterious load in mutators, non-mutators were always competed against non-mutators carrying the opposite fluorescent marker while mutators were always competed against mutators. Fitness values of evolved populations reported in Figure 3 and Extended Data Figure 4 were normalized by the relative fitness of their ancestral strain.

**Extended Data Fig. 4.**
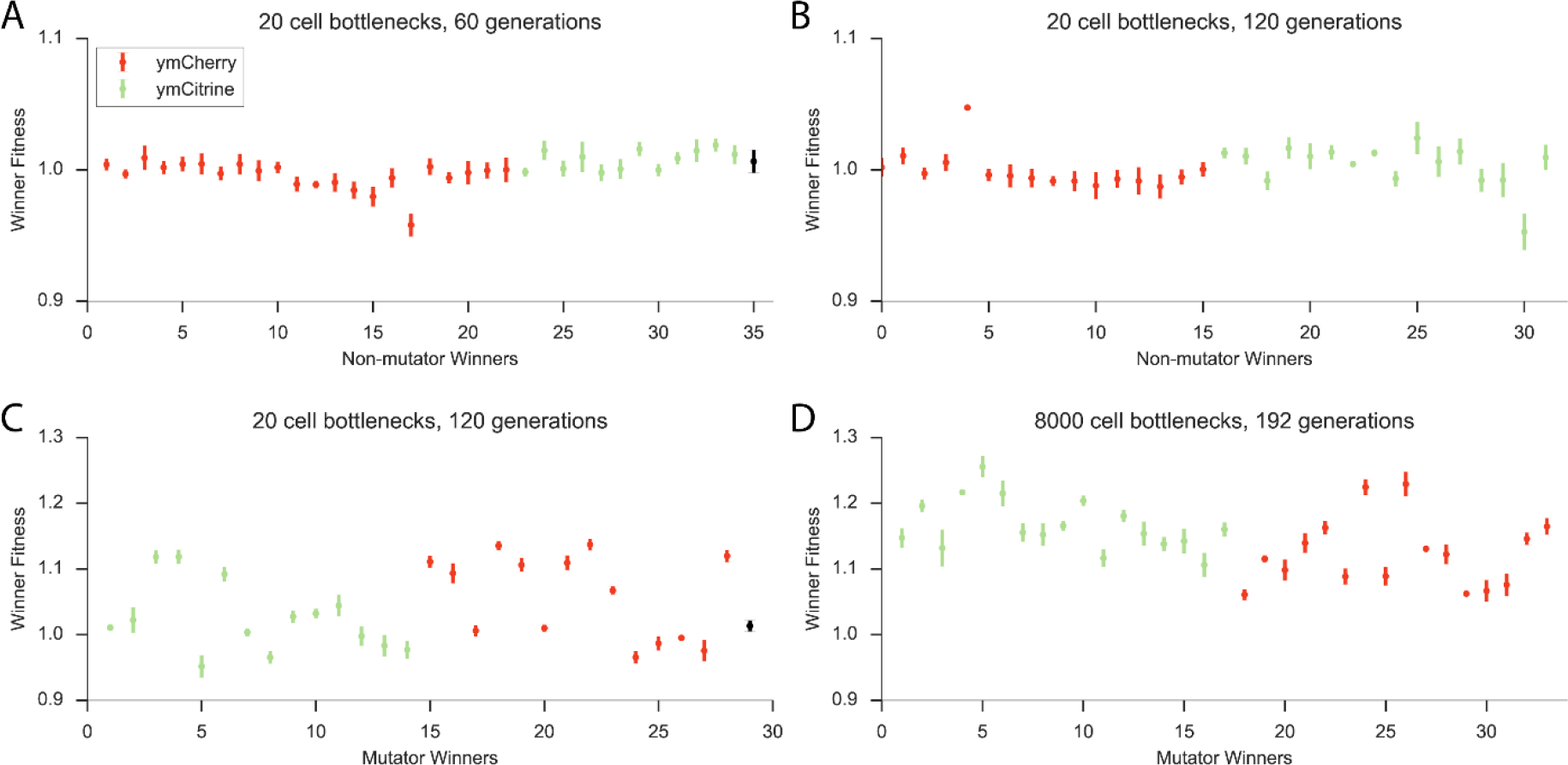
Fitness values of competition winners. Each point represents an average of 5 replicates, error bars ±95% CI. In black, relative fitness of ancestral ymCherry-labeled strain (non-mutator in panel A and mutator in panel C) against the ymCitrine-labeled strain.

### Statistical Analysis

A two-sided binomial test^41^ was used to assess whether realized fixation probability in our experiments was significantly different from the neutral expectation (given by the starting frequency). A two-sided paired t-test^41^ was used to compare mean mutator frequency at the end of the experiment to mean mutator frequency at the beginning. A two-sided unpaired t-test^41^ was

### Data Availability

Frequency trajectories of our competitions in Figure 2 are contained in Supplementary Data 1. Relative fitness values of all the competition winners in Figure 3 are contained in Supplementary Data 2.

### Code Availability

Simulation code is available at https://github.com/yraynes/Sign-Inversion.

